# Immunorecognition of *Streptococcus mutans* secreted proteins protects against caries by limiting tooth adhesion

**DOI:** 10.1101/2023.06.21.545893

**Authors:** Omer Bender, Jessica Khoury, Gal Hirsch, Evgeny Weinberg, Naor Sagy, Shani Buller, Shiri Lapides-Levy, Sigalit Blumer, Daniel Z. Bar

## Abstract

Childhood caries, a chronic disease affecting 60–90% of children in industrialized countries, results in lesions in primary and permanent dentition, leading to hospitalizations, emergency room visits, high treatment costs, and loss of school days. It diminishes the child’s ability to learn and increases the risk of caries in adulthood. Despite multiple risk factors for caries, significant interpersonal variability unaccounted for by these factors exists. The immune system generates a personal antibody repertoire that helps maintain a balanced and healthy oral microbiome. *Streptococcus mutans* is a key bacterium in caries development. Utilizing mass-spectrometry, we examined which *S. mutans* proteins are identified by antibodies of children with and without caries and identified a core set of proteins recognizable by the immune system of most individuals. This set was enriched with proteins enabling bacterial adhesion. To study the physiological relevance of these findings, we tested the ability of saliva to prevent *S. mutans* adherence to tooth surfaces. Saliva from caries-free individuals, but not children with caries, was found to hinder the binding of *S. mutans* to teeth. These findings delineate the *S. mutans* proteome targeted by the immune system and suggest that the inhibition of bacterial adherence to teeth is a primary mechanism employed by the immune system to maintain oral balance and prevent caries formation. These discoveries offer fresh insights into the immune system’s role in preserving oral health and preventing caries development. Targeting *S. mutans* proteins implicated in bacterial adhesion could be a promising strategy for preventing childhood caries.

## Introduction

### Dental caries

Dental caries is a worldwide, multifactorial disease caused by an imbalanced oral environment and correlated, among others, with cariogenic diet, lack of oral hygiene and thriving of oral cario-pathogens, genetic factors, and socio-economic factors ^1,2^. Caries is one of the most prevalent chronic conditions among children, impacting 60-90% of youngsters in developed countries^3^.

The onset of caries increases the likelihood of experiencing dental decay in later years, which may necessitate complex dental treatments ^4,5^. The consequences of childhood caries often include a higher risk of new carious lesions in both the primary and permanent dentitions, hospitalizations and emergency room visits, high treatment costs, loss of school days, diminished ability to learn, and impaired oral health-related quality of life. Moreover, families are affected through treatment expenses, work absences, and parental feelings of guilt ^6–8^.

### Caries and Bacteria

The oral cavity is considered one of the most densely populated sites within the human body, harboring over 700 bacterial species ^9,10^. Aerobe and anaerobe bacteria together form multi species communities known as biofilm. As soon as a tooth surface erupts or is cleaned, salivary proteins and glycoproteins are absorbed, forming a conditioning film termed the acquired pellicle ^9,10^. Bacterial components, such as glycosyltransferases and glucan binding proteins, form bonds with the acquired pellicle ^11,12^. This formation anchors the biofilm to the tooth surface, resists changes in the oral environment, and allows the proliferation of these bacterial communities.

The emergence of dental caries involves interactions between the tooth structure, the microbial biofilm, and sugars, and is influenced by the saliva content and genetic factors. Dental caries typically initiate at the enamel surface as a result of a process in which the crystalline mineral structure of the tooth is demineralized by organic acids produced by the bacterial biofilm. The loss of minerals leads to cavity formation and further progression of bacterial acid secretion within this protected environment, thus creating a caries lesion ^1,2^. The ecological changes in the oral cavity lead to the development of a biofilm that is more cariogenic. This is predominantly, but not exclusively, driven by the bacterium *Streptococcus mutans. S. mutans* is strongly associated with dental caries, mainly because it holds many caries-contributing mechanisms^13^.

One of those mechanisms, tightly connected to the disease pathogenesis, is the ability of *S. mutans* to adhere to solid surfaces like the tooth enamel. Through the mechanism of adhesion to a solid surface, *S. mutans* are capable of colonizing the oral cavity and forming bacterial biofilm. Several cell-surface and secreted proteins are important for *S. mutans* adhesion.

These include antigen I/II (AgI/II); glucosyltransferases (GtfB, GtfC, GtfD) and glucan-binding proteins (GbpA, GbpB, GbpC, GbpD)^14–16^.

### The Oral Immune System

The oral mucosal immune system is composed of mucus layers, epithelial cells, lymphoid tissues, and immune molecules ^17^. The mucosal immune system maintains a balance between tolerance to commensal bacteria and immunity to pathogens mainly through SIgA, being the predominant antibody isotype in the oral cavity^18^. SIgA, a subtype of IgA, differs from it by an additional secretory component. Its main function is to bind to bacterial surfaces and virulent secreted factors, preventing their activity. These antibodies are formed from V, D, and J gene segments which undergo DNA rearrangement (referred as V(D)J recombination), resulting in a diverse set of unique antigen receptors ^19^. The extensive array of combinations allows for an individualized immune response against *S. mutans*, which could be a factor in the variability observed in DMFT (Decayed, Missing due to caries, and Filled Teeth) scores among children and adolescents. The immune response to *S. mutans* is dependent on caries status and age, with age-related affinity maturation potentially facilitating the production of more effective antibodies ^20^. Favorable conditions brought about by the immune system might promote a healthy oral microbiome by limiting or preventing the colonization of cariogenic bacteria on tooth surfaces.

Although antibodies to individual proteins have been investigated in relation to their role in caries, a comprehensive, hypothesis-free mapping of *S. mutans* antigens has yet to be conducted.

In the present study, we measured the protein-specific immune response to the cariogenic bacterium *S. mutans* in the whole saliva of a cohort of 34 children between the ages of 6 - 12. This cohort included caries-free participants (DMFT = 0; n = 17) and participants with high levels of cariogenic activity (DMFT≥5; n = 17). Using mass spectrometry, we identified proteins aiding in bacterial adhesion as the prime target of the immune system. These key findings were validated using the enzyme-linked immunosorbent assay (ELISA) on additional participants (N =170) of similar ages. Furthermore, our results demonstrated that saliva from caries-free individuals prevents *S. mutans* from binding to tooth surfaces. These results provide a comprehensive map of the *S. mutans* proteome targeted by the immune system, suggesting that inhibition of tooth attachment is a primary mechanism employed by the immune system to maintain oral balance and prevent caries.

## Results

### Characterizing *S*. *mutans* Proteins Recognized by the Immune System

To identify the proteins recognized by the immune system and to determine their correlation with the dental caries status of each participant, we isolated antibodies from saliva and characterized the *S. mutans* antigens that they target (Fig. 1). We compared two groups of children from similar environments and socioeconomic backgrounds: a caries-free group (n=17) and a high DMFT group (n=17) (Fig 1A). The entire spectrum of antibodies was collected from each saliva sample using Protein-L beads. These were then used to bind antigens from a whole lysate of *S. mutans*. Captured proteins were digested, multiplexed and analyzed using mass spectrometry (Fig. 1B). Each TMT multiplexed proteomics run included the pull-down of four participants from each group, a negative control group that omitted the antibodies from the full antigen precipitation, and a positive control group of diluted whole *S. mutans* lysate.

**Figure 1:**
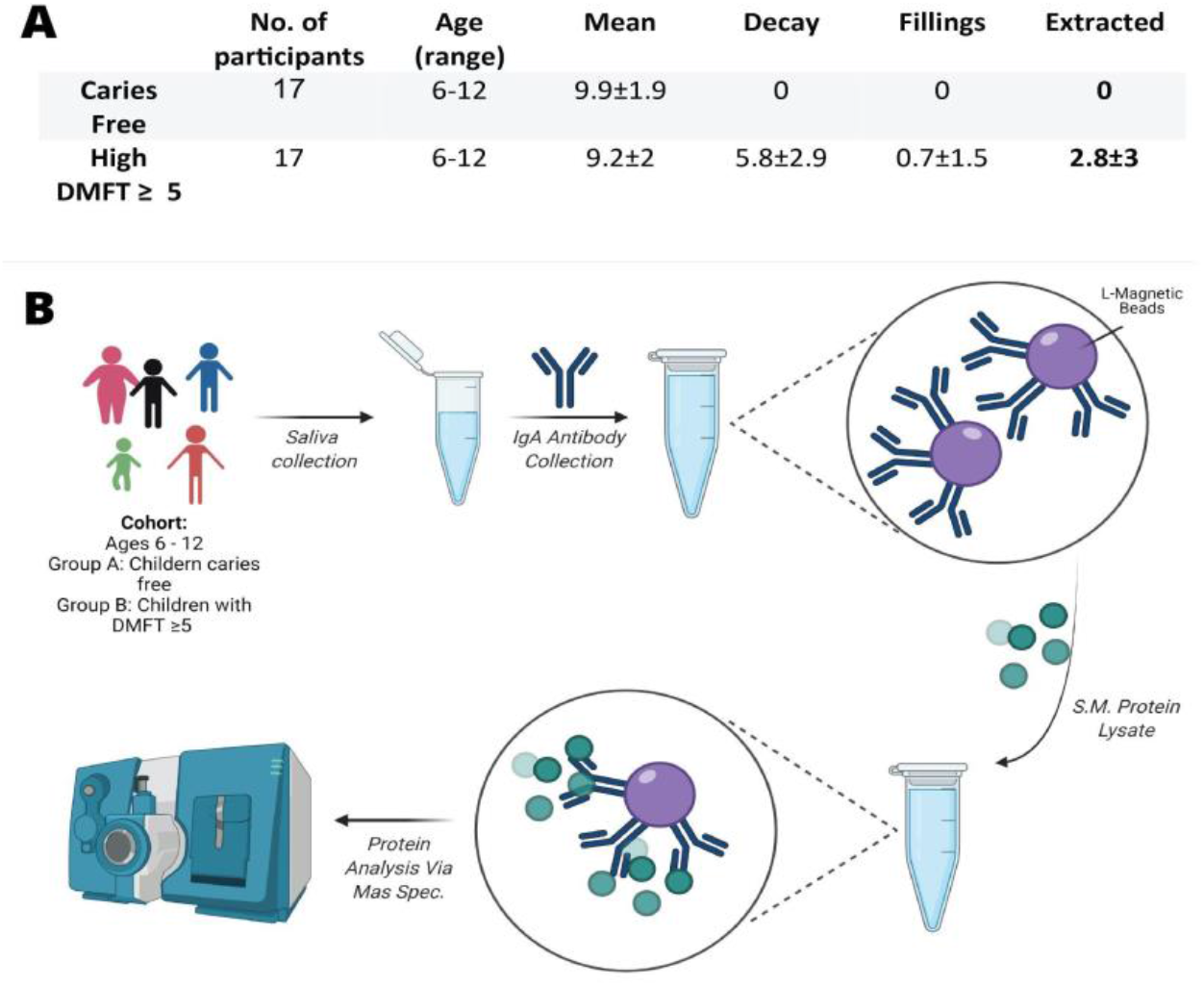
**A**. Statistical data of the cohorts participants. Values are mean values with their standard deviation. **B**. Schematic Overview: Whole saliva obtained from 34 children patients from the Goldschleger School of Dental Medicine after consigned consent from legal guardians, saliva antibodies are captured on protein L affinity beads. Solubilized antigens (proteins) in an S.mutans lysate bind the antibodies anchored to the beads, while non-specific and weak binders are washed away. After elution, proteins are digested, TMT labeled, multiplexed, cleaned and sent for identification by mass-spec analysis.

Of the 1953 known *S. mutans* proteins, 44 proteins were identified by the immune system of at least some of the participants. The network of these protein interactions is available at Supplementary Data. Of these, 10 proteins were identified by antibodies of most participants. For instance, in one MS run with eight participants (four from the caries-free group and four from the high DMFT group), we observed that nine out of ten proteins passed our detection threshold. Protein binding appeared stronger in the high DMFT group compared to the caries-free group (Fig. 2A), but this trend was not statistically significant for individual proteins.

**Figure 2:**
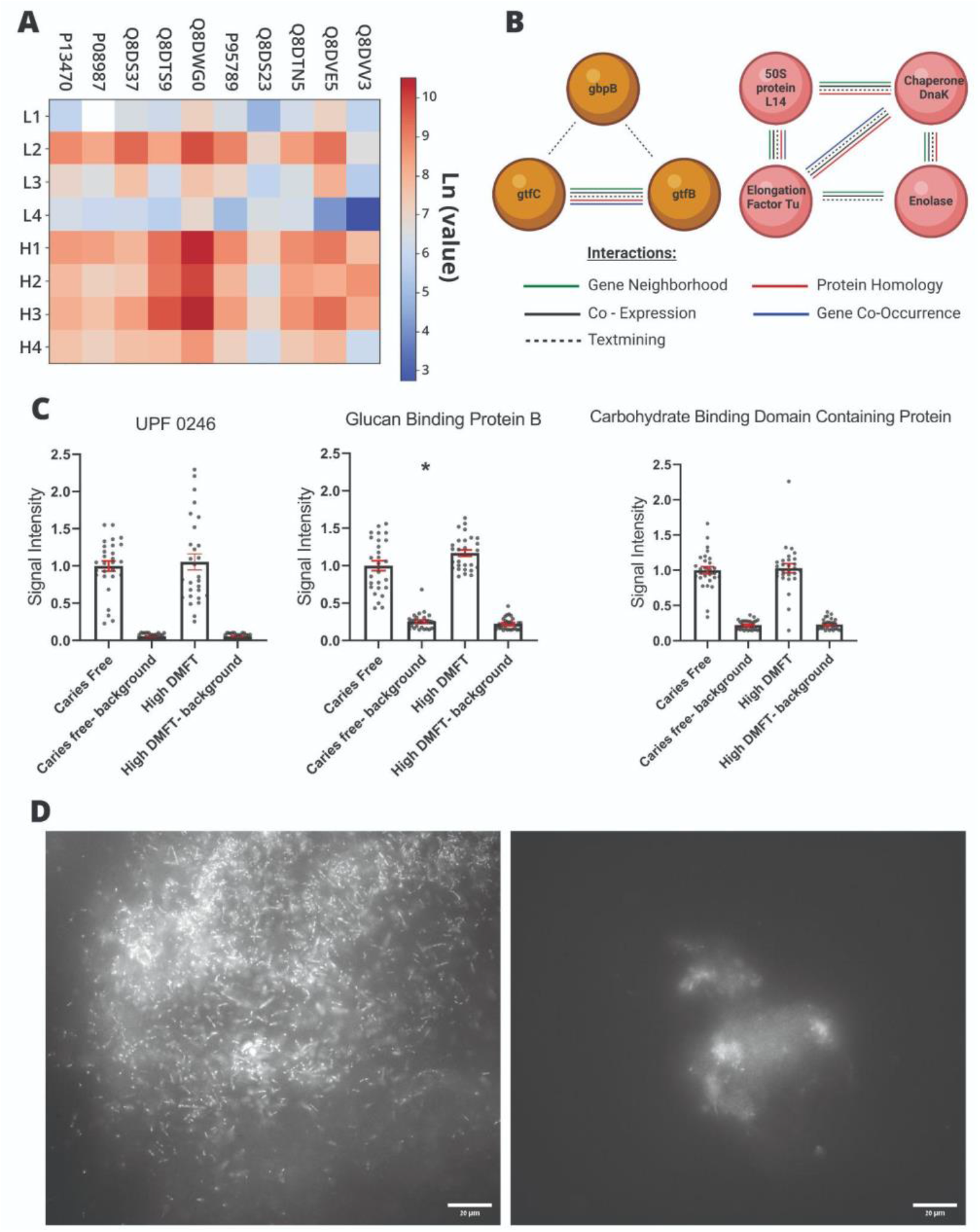
**A**. Heat map of results, Ln (value) of relative intensity from a TMT 10 plex run with 4 samples of caries free children (L1-4) and children with DMFT ≥ 5 (H1-4), positive and negative controls are not shown. **B**. Protein interactions of the proteins found in most participants in more than one TMT 10-Plex MS run using the String database (www.string-db.org). **C**. Sandwich ELISA results of 3 selected proteins representing the differences in signal intensity of each experiment to background noise and comparing the signal intensity of low and high DMFT groups to the different S. mutans antigens at 450 nm. A significant signal above the background was recorded in all three cases. UPF0246 - No statistical difference between caries free and high DMFT groups. GbpB - Statistically higher signal detected in the high DMFT group (P<0.05). Q8DWI5 - No statistical difference between the two groups. **D**. Representative image of bacterial attachment and growth on deciduous teeth. a. Bacterial tooth attachment using saliva of a child with high DMFT values. We can observe a high density of colonies of S. mutans dyed with DAPI, on the surface of a deciduous tooth. B. Bacterial tooth attachment using saliva of a caries free child. Images were acquired on a LSM 900 microscope (Zeiss) using a 63X oil lens.

Several of the 10 core proteins identified (Table 1) are involved in the attachment of *S. mutans* to hard surfaces. For example, Glucosyltransferases B and C, which catalyze the synthesis of insoluble glucans from sucrose, mediate the firm adherence of cells to tooth surfaces and support the formation of the biofilm. Meanwhile, glucan-binding proteins (GbpA, GbpB) modulate cell membrane stability and stabilize adherence to extracellular glucans. These proteins, which are known to contribute to caries, elicit an immunogenic response ^21,22^. Among the ten identified genes, seven had known relationships and formed two distinct clusters (Fig. 2B). These clusters were characterized by a common role in hard surface adhesion, high genetic similarity (homology), chromosomal proximity, and/or co-expression.

**Table 1:** Proteins found in most participants in more than one TMT 10-Plex MS run.

| Uniprot ID | Name | Known and putative functions |
| --- | --- | --- |
| P13470 | GtfC/ Glucosyltransferase-SI | Synthesizes both water-insoluble (alpha 1,3-linked glucose and some 1,6 linkages) and water-soluble glucans (alpha 1,6-glucose). Secreted. |
| P08987 | GtfB/C/Glucosyltransferase-I | Synthesizes water-insoluble glucans. Secreted. |
| Q8DWM3 | GbpB | Putative secreted antigen GbpB/SagA putative peptidoglycan hydrolase. |
| Q8DTS9 | Enolase | Binds plasminogen and human salivary mucin MG2. May enable bacteria to acquire surface-associated proteolytic activity and aid in dissemination through oral tissues. |
| Q8DVE5 | Tagatose 1,6-diphosphate aldolase 2 | Carbohydrate metabolism |
| O06942 | Chaperone protein DnaK | Maintains protein homeostasis during cellular stress through two opposing mechanisms: protein refolding and degradation. |
| Q8DS23 | 50S ribosomal protein L14 | Ribosomal protein. |
| P72483 | Elongation factor Tu | Promotes the GTP-dependent binding of aminoacyl-tRNA to the A-site of ribosomes during protein biosynthesis. Also plays a role in the regulation of autophagy and innate immunity. |
| Q8DRY6 | Uncharacterized Protein | Uncharacterized protein. |
| Q8DWI5 | Carbohydrate-binding domain-containing protein | Uncharacterized protein. |

### Validating Differential Immunological Response to *S*. *mutans*

We selected five proteins for further validation of the mass spectrometry findings using an independent method. The selected proteins included two that are known to be important, GBPC and GBPB (Table 1), two proteins of uncharacterized function, UPF0246 and Q8DWI5, and Elongation Factor Tu. We successfully cloned, expressed, and purified four of the five proteins from *E. coli*. However, despite multiple attempts, we could not express GBPC. We hypothesized that it was toxic to *E. coli*. Afterwards, we carried out a sandwich ELISA to test the salivary immune response of the low and high DMFT groups to these selected *S. mutans* antigens. Nearly all tested participants exhibited a robust immune response to three out of the four purified proteins, but none responded to the Elongation Factor Tu.

The sandwich ELISA plates incubated with the three remaining proteins showed significant signals above background in both groups being examined (Fig. 2C). Consistent with the mass spectrometry data and published literature ^23^, we observed strong binding for GbpB in all samples. However, the immune systems of the high DMFT patients showed a preference for binding to GbpB (p = 0.005), suggesting that GbpB is a principal target of the immune system (Fig. 2C) in its efforts to maintain local homeostasis.

In line with the mass spectrometry data, Q8DWI5 and UPF0246 also generated robust signals, indicating the presence of protein-specific antibodies. However, we observed no difference between the caries-free and high DMFT groups.

### Saliva inhibits *S*. *mutans* plaque formation on teeth in a DMFT-dependent manner

One interpretation of the composition of recognized antigens is that the immune system prevents or delays the attachment of *S. mutans* to the tooth surface. To test our hypothesis, we examined the binding-inhibition capacity of saliva from children with high or low DMFT scores (Table 2). Specifically, we incubated *S. mutans* bacteria with saliva in a nutrient-rich medium. We then briefly allowed the bacteria to bind to extracted deciduous teeth. We allowed the bound bacteria to grow overnight, stained them for DNA, and imaged them the next day. Saliva from caries-free children significantly inhibited tooth attachment compared to that from children with high DMFT scores (P=0.0136; Fisher’s exact test, Fig. 2D). These results, from both the sandwich ELISA and plaque formation tests, support the hypothesis that the immune system prevents *S. mutans* from colonizing tooth surfaces by using secreted antibodies, possibly allowing the attachment of other bacteria to the teeth.

**Table 2:** S. mutans tooth attachment scores in the presence of saliva from caries-free and high DMFT donors.

| | Caries free | DMFT $\geq 1$ | Total |
| --- | --- | --- | --- |
| **Classified high** | 17 | 45 | 62 |
| **Classified low** | 18 | 15 | 33 |
| **Total** | 35 | 60 | 95 |

## Discussion

### Immunorecognition of *S*. *mutans*

Dental caries, especially among children, is one of the most prevalent diseases worldwide ^4,24^. Secretory antibodies in saliva and gingival crevicular fluid serve as a primary defense against pathogenic bacteria, playing a critical role in maintaining a balanced and beneficial oral microbiome ^25,26^. Research has explored the immunogenicity of *S. mutans* and its relationship to caries, identifying specific virulence factors. However, unbiased data on the antibody-specific targets of *S. mutans* and the effectiveness of these antibodies in neutralizing the bacterium and its virulence factors remain scarce. By immobilizing secreted antibodies from the oral cavity, and using them for immunoprecipitation of their targets, we have identified 44 different proteins recognized by different individuals’ immune systems. Using mass spectrometry, we found that most of the cohort identified 10 *S. mutans* proteins. These shared proteins form the basis of the common immunological response to *S. mutans*. This distribution aligns with protein and epitope profiling for other diseases and pathogens, where a core of proteins and epitopes is shared, while additional targets are recognized by the immune system of a few individuals ^27–29^. These findings reinforce much of the existing literature by providing evidence of immunorecognition of several secreted proteins with crucial roles in adhesion to hard surfaces and extracellular matrix organization.

### Efficacy of virulence factors neutralization

To validate the mass spectrometer results independently, we constructed plasmids expressing our proteins of interest: Elongation factor Tu, UPF0246, GbpB, GbpC, and Q8DWI5. We were able to express and purify these proteins in *E. coli*, except for GbpC, possibly due to its toxic effects on the bacteria. Finally, we performed S-ELISA, using saliva samples from children with high and low DMFT scores as the source of primary antibodies.

As stated previously, significant statistical differences between the two groups were only observed when incubating sandwich ELISA plates with the GbpB proteins. These results are consistent with previous findings ^23^. The high DMFT group bound to GbpB stronger than the low DMFT group, indicating a greater presence of antibodies targeting GbpB in high DMFT score patients. These results may be attributed to the continuous exposure of high DMFT patients to *S. mutans*, leading to the generation of more antibodies against GbpB.

In contrast, the antibodies of children from both the low and high DMFT groups bind equally to Q8DWI5 and UPF0246 proteins. These results do not exclude the possibility of a differential immunogenic response. Our purified proteins might exhibit differences in post-translational modifications, given that they were purified from an *E. coli* expression system. These modifications can generate distinct epitopes for the same antigen, eliciting different immune responses when expressed by a different bacterium ^30^. Additionally, these results did not address the specific epitopes on each protein that may also differ between the two groups. Thus, these results should not be used to exclude the involvement of Q8DWI5 and UPF0246 in caries or that of antibodies against them in caries protection. Instead, further investigations are required to assess the reactivity to these proteins and their involvement in caries formation.

In all four repeats of the experiment, Elongation Factor Tu did not generate a detectable signal. One possibility is that this highly abundant internal bacterial protein was a false positive identification by our mass-spectrometry experiments. However, secreted variants with roles in the extracellular matrix have been reported in other organisms ^31^ and epitope mapping in other streptococci identified it as highly immunogenic ^32^. Therefore, it is possible that the different post-translational modifications in the *E. coli* resulted in a non-immunogenic version of an otherwise immunogenic protein.

### Biofilm formation proteins

The acquired pellicle initiates the formation of biofilm on the hard tooth surface. Several oral microbiota, including *S. mutans*, adhere to the pellicle. *S. mutans* adherence to the pellicle is promoted by high concentrations of sucrose. One major class of proteins found is biofilm formation proteins. These include Glucosyl-Transferase (Gtf) B, which primarily synthesizes insoluble glucans, GtfC, which primarily synthesizes soluble glucans, and Glucan-Binding-Protein (Gbp) B, GbpC, and Enolase, which assist *S. mutans* and other bacteria in adhering to tooth surfaces. Interestingly, GbpB is not only considered important for the cariogenicity of *S. mutans*, but mucosal immunization in animal models has been found to induce a protective effect against caries ^33,34,35^. Glucosyl transferases B and C are recognized as major cariogenic virulence factors of *S. mutans*. One study demonstrated an increased amount of Glucosyltransferase isoforms in in situ pellicles of children with high DMFT scores compared to those with low DMFT scores ^36^. Furthermore, a correlation between high levels of GtfB and high DMFT levels has been previously established ^37^. Interestingly, our findings show an increase in the recognition of the adaptive immune system against these proteins (Fig. 2C). This suggests a faulty mechanism in the immune system’s ability to neutralize these proteins, thereby promoting, or failing to inhibit, *S. mutans* adherence to tooth surfaces (Fig. 2D).

### Novel immune-reactive proteins

We have identified multiple novel proteins as targets of the immune system. These include uncharacterized proteins, some of which are targeted by the majority of individuals in the study. For example, Q8DRY6, or protein SMU 2070, has no known activity, while Q8DWI5, or Uncharacterized protein UP000002512, contains a Carbohydrate-binding domain. ELISA measurements have detected no difference in binding of Q8DWI5 by antibodies of children with low and high DMFT. Another interesting protein identified in the mass-spectrometry analysis is the Chaperone protein DnaK. DnaK is a heat shock protein that is involved in the regulation of protein synthesis. Elevated levels of this protein were identified in *S. mutans* in response to acidic and heat stress ^38,39^, suggesting that higher concentrations of these proteins contribute to its durability. It has also been shown that knock-out of this protein lowers the virulence capabilities of *S. mutans* by inhibiting biofilm formation and acid tolerance ^40^.

We suggest that while children with high DMFT scores generate more antibodies targeting these proteins, these antibodies are less efficient at preventing the bacteria from attaching to teeth, and therefore have a more limited capacity of maintaining oral balance.

## Conclusions

The present study contributes to the understanding of the immunoreactive targets of *S. mutans*, a key pathogen responsible for dental caries in children and adolescents. Our findings, which are consistent with previous literature, suggest that biofilm proteins are among the major virulent risk factors contributing to the development of caries ^2,14^. The inhibitory properties of saliva from the caries-free group, coupled with the higher quantity of antibodies in the high DMFT score group identified by mass spectrometry, suggest a deficiency in the immune system’s ability to neutralize *S. mutans* or its virulence factors. Notably, our data underscore the potential for targeting these proteins to mitigate caries risk. Furthermore, monitoring antibody levels to these targets could aid in identifying at-risk individuals and implementing preventative measures at an earlier stage. Overall, these findings have significant implications for the prevention and management of dental caries, a prevalent and costly public health issue. Future work to identify binding sites through peptide mapping could help pinpoint more specific targets, thereby informing potential therapeutic interventions and vaccine development.

## Materials and Methods

### Ethical Statement

This work was approved by the Ethics committee of Tel-Aviv University (Proposal no. 0000743-2).

### Sample Collection

Saliva samples were obtained from children between the ages of 6-12 at the Goldschleger school of dental medicine clinics in Tel Aviv University, after informed consent and legal guardian signed approval.

Saliva was collected by spitting unstimulated saliva into a 50 mL tube. The whole saliva was centrifuged at 12,000 RCF for 7 minutes. Supernatant, containing the soluble antibodies, was separated and stored at -80°C until use.

### Antigen Preparation

*Streptococcus mutans* strain ATCC UA159 was obtained from Prof. Gilad Bachrach (The Hebrew University of Jerusalem, Israel). We confirmed it as *S. mutan*s through targeted PCR using the primers TCGCGAAAAAGATAAACAAACA and GCCCCTTCACAGTTGGTTAG, sequencing of the resulting product, and protein mass-spectrometry. Bacteria were cultured according to ATCC guidelines (*Streptococcus mutans* Clarke). For antigen preparation, we cultivated the bacteria under anaerobic conditions in a Brain Heart Infusion (BHI) medium for 48 hours. Cells were centrifuged for 30 minutes at 2000 RFC using an Eppendorf 5910R centrifuge, medium discarded, cells were washed with PBS and pellet resuspended in RIPA lysis buffer, followed by sonication for 10 minutes at 20% power (OMNI International - Sonic Ruptor 400). Following centrifugation at 4500 RCF for 30 minutes, we discarded the pellet. Protein concentration was measured using nanodrop OneC (Thermo-Fisher) 205 nm method. Proteins were aliquoted at 1 mG/mL and stored at -80°C until use.

### Study Design

Reactivity of saliva antibodies against *S. mutans* were evaluated in two different groups of children between the ages 6 – 12: caries free (DMFT=0; n=17) and high-DMFT (DMFT≥5; n=17). Antibody-bound proteins were identified using mass spectrometry. Selected proteins were further validated using ELISA on additional saliva samples.

### MS Sample preparations and analysis

Briefly, saliva antibodies from 34 participants were captured on protein L affinity beads. Solubilized *S. mutans* antigens (proteins) were allowed to bind the antibodies anchored to the beads. After washes, eluted proteins were digested, TMT labeled, multiplexed, cleaned and sent for identification by mass spectrometry analysis. From the identified proteins, we filtered out proteins with less than 3 peptides. To account for non-specific binding, a ratio of the relative intensity of the positive control (diluted protein lysate) to negative control (full protocol with antibodies omitted) was calculated.

### Plasmid construction expressing proteins of interest and ELISA assay

Five plasmids were engineered to express the following genes: Elongation factor Tu (P72483 Uniprot ID), UPF0246, GbpC (Q8DFT1 Uniprot ID), GbpB (Q8DWM3 Uniprot ID), Carbohydrate-binding domain-containing protein (Q8DWI5 Uniprot ID). The construction of these plasmids was based on the pDL278_P23-DsRed-Express2 vector’s backbone ^41^. pDL278 vector was selected since it replicates and stays stable in various oral streptococci, besides being suitable for cloning and expression in *E. coli*. This vector incorporates the P23 promoter, allowing constitutive expression of genes, and a gene encoding for DsRed-Express2 protein, which is a red fluorescent protein. For plasmids construction, first the protein sequences were downloaded from uniprot, and the codon-optimized genes were cloned into the pDL278_P23-DsRed-Express2 vector, replacing the DNA sequence of the gene expressing DsRed-Express2. To the C-terminal of the gene sequences, 6xHistag (HHHHHH peptide sequence) and Flag tag (DYKDDDDK peptide sequence) were added. HIS tag for easing purification and a flag tag for confident detection. Illustration of the structure of the plasmid expressing UPF0246 protein: The other plasmids of proteins of interest have similar structure but differ at the gene sequence region.

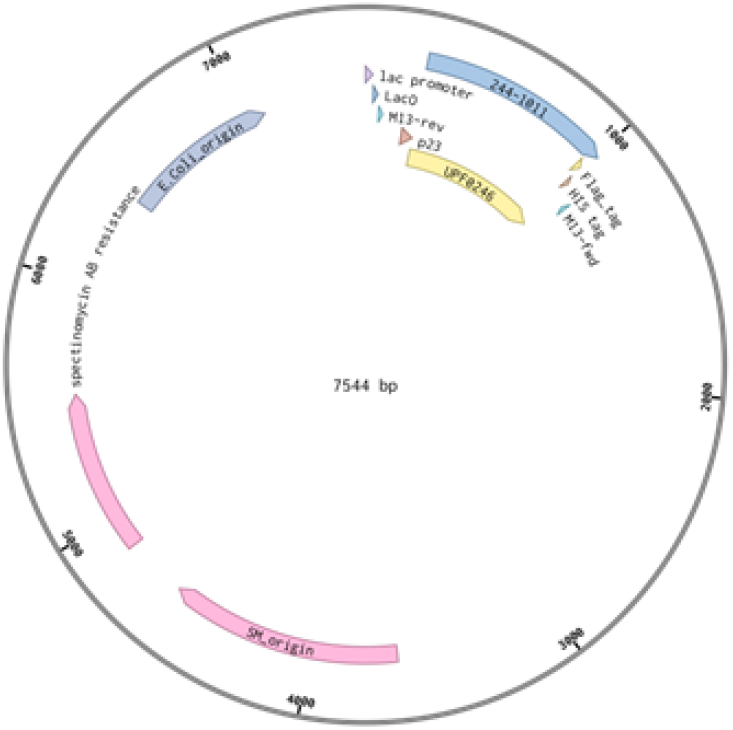

### Bacterial protein expression

The plasmids were transformed into *E. coli* strain BL21 for protein expression. One bacterial colony was selected from the petri dish and grown overnight in LB medium with 50 μg/mL of spectinomycin at 37 °C and 180 x g speed. The following day, the cells were collected and pelted by centrifugation at 4500 x g for 45 min at 4 °C, and pellets were frozen and stored at - 80 °C.

### Protein purification by Nickel beads

The frozen pellets were lysed by suspending them with 20 mL lysis buffer made of 50mM phosphate buffer, 150 mM NaCl, 10 mM imidazole, one pill inhibitor protease cocktail (Mercury, REF. 11836170001), tween 20 0.5%, and 0.1 mM EDTA. After suspension, cells were disrupted by two cycles of sonication (OMNI International - Sonic Ruptor 400) cycles on ice, 8 minutes each (40 power level). The lysates were centrifuged at 4 °C for 30 min at max speed (Eppendorf 5910R, 20133 rcf). The supernatants, containing the proteins of interest, were collected and diluted with a 10 mL of equilibration buffer and nickel (Ni) beads. The equilibration buffer was composed of 50 mM phosphate buffer, 150 mM NaCl and 10 mM imidazole. The supernatants along with the Ni beads were placed on a rotator at 4 °C for 4 hours of incubation, providing enough time for the Ni beads to bind to His-tagged proteins. Afterwards, they were centrifuged at 4 °C for 5 min at 500 x g. The supernatants were discarded and beads were resuspended in a 10 mL wash buffer, constituting 50 mM phosphate buffer, 150 mM NaCl, and 40 mM imidazole. The resuspensions were stirred at 4 °C for 15 minutes and later were centrifuged at 4 °C for 15 min at 500 x g. Again, the supernatants were aspirated and pellets were eluted by 5 mL elution buffer, made of equilibration buffer with 300 mM of imidazole. Then, the resuspensions were stirred at 4 °C for 15 minutes and finally were centrifuged at 4 °C for 15 min at 500 x g. The supernatants were collected and their protein concentration was measured with the aid of nanodrop OneC (Thermo-Fisher) by using the A280 method. The purification yields for elongation factor TU, UPF0246, glucan binding protein B and carbohydrate-binding domain-containing protein were 0.057 mg, 0.165 mg and 0.057 mg and 0.033 mg per 1 liter, respectively. By contrast, glucan binding protein C could not be purified either because *E. coli* BL21 strains failed to produce colonies or failed to produce colonies which express the protein of interest. Trials with additional E. coli strains also failed. As a consequence, we concluded that the constitutive expression of this protein is toxic to *E. coli*.

Supernatant subsamples were used for expression validation via SDS-PAGE gels and western blots. The protein-containing solutions were diluted by 50% glycerol and then they were stored at -80 °C.

### ELISA

The ELISA assay was performed using a compatible PVC clear high bind ELISA microtiter 96-well plate. First, 100 μl per well of saliva samples was used for coating collected from 2 groups of patients in ages between 6-12 years old; low DMFT (DMFT=0) patients and high DMFT (DMFT≥5). Prior to the binding step, the saliva samples were diluted 10 times in PBST 0.05% + 1% BSA. The plate was placed on a rotator and incubated overnight at 4 °C. The next day, it was washed three times with 200 μl of PBS-T 0.05%. 200 μl blocking solution containing PBST 0.05% and 5% BSA were added to each well, and the plate was incubated for 4 hours at RT on a rotator to prevent non-specific bindings. After incubation, the plate was inverted, the solution was flicked out, and washing steps were repeated. Binding was done with 100 μl of the purified proteins, diluted to a desired concentration at a final volume of 100 μl per well. The plate was placed on a rotator and incubated for 40 minutes at RT. This was followed by two fast washes with PBST 0.05% and three long washes with PBST 0.05% + 1% BSA, each duration of wash was about 10 minutes waiting time on a rotator at RT. Afterwards, 100 μl of anti-flag HRP-conjugated antibody (1 μl antibody diluted into 5 mL PBST 0.05%) were pipetted to relevant wells and incubated for 30 minutes at RT. The last described washing steps were repeated and eventually 100 μL of TMB was added to each well and the plate was incubated at RT (4 – 30 min) for colour-reaction development. Finally, if colour develops, 50 μL of 0.5 M H2SO4 was added to each well to stop the enzymatic reaction. Plate absorbance was immediately measured at 450 nm absorbances using a plate reader (Synergy H1, Biotek).

### *S*. *mutans* tooth adherence assay

The ability of saliva in preventing *S. mutans* attachment to teeth was measured by brief incubation of cleaned teeth in BHI containing *S. mutans* in the presence of saliva from individuals with high (n = 10) and low (n = 10) DMFT scores. Detailed protocol available at Supplementary Data. Teeth were washed and moved to fresh wells with growth medium but without bacteria, and adherent bacteria were allowed to grow overnight. The teeth were examined under LSM 900 microscope (Zeiss) using a 63X oil lens and 3-5 images were taken from each tooth. The images were scored by two blinded evaluators, asked to assign scores as follows: 1 = no bacterial growth, 2 = low bacterial growth and 3 = high bacterial growth. The overall score for each tooth sample was calculated. Fisher’s exact test was performed on the evaluators’ responses.

### Bioinformatics analysis

The six raw data files were searched against the *Streptococcus Mutans* UA159 proteome (uniprot #UP000002512) using MaxQuant V2.1.0.0 software (Max Planck institute). The minimum peptide count per protein was set to 3, with FDR set to 10% per peptide. Reverse sequences were used for decoy search and contaminant sequences were included in the search. Only proteins with a relative intensity ratio of positive control to negative control greater than 1 were included in our data analysis.

### Statistics

Demographics and clinical characteristics were presented as mean with standard deviations (Mean ± SD). Unless otherwise noted, p-values were calculated using a two-tailed T-test. Bacterial adhesion experiments were performed twice and ELISA experiments were repeated four times.

## Data availability

Generated data is available at ProteomeXChange via PRIDE repository; Identifier PXD043103.

## Acknowledgments

We thank the members of the Bar lab for their critical reading of the manuscript, useful suggestions, and comments. We thank the Israeli Science Foundation for funding (D.B. 632/20). We thank the Smoler National Proteomics Centre at the Technion, Israel, for the operation of the mass spectrometer and technical assistance in sample preparation. We thank Ariel Pokhojaev for his assistance in formatting and editing the figures.

O.B. contributed to experimental conception, design, data acquisition and interpretation, drafted the manuscript. J.K. contributed data acquisition and interpretation, and drafted the manuscript. G.H. contributed data acquisition and interpretation. E.W. contributed to experimental design and data analysis. N.S. contributed data acquisition and interpretation.

S.B. contributed data acquisition and interpretation. S.L.L. contributed data acquisition and interpretation. S.B. contributed to funding, conception, design, data interpretation, drafted and critically revised the manuscript. D.Z.B.contributed to funding, conception, design, data interpretation, performed all statistical analyses, drafted and critically revised the manuscript.

