## Supplementary Data for "Immunorecognition of *Streptococcus mutans* secreted proteins protects against caries by limiting tooth adhesion"

**Saliva antibody immobilisation:**

1. Add 100 µL of saliva after spin-down to 40 µL of L - Magnetic bead in a 1.5 mL tube.
2. Add 860 µL of PBS with 0.05% Triton-X 100, leave overnight in a 3D orbital shaker at 4 C.
3. Put the tube in a magnetic holder, remove the supernatant, add 500 µL PBS with 0.05% Triton-X 100 and put in a 3D orbital shaker for 5 minutes.
4. Repeat the washes 3 times.

**Antibody - Antigen binding:**

5. Add 500 uG of protein lysate in 100 uL to the sample and add PBS with 0.1% SDS and 0.1% Triton-X 100 up to 1ml.
6. Leave overnight in a 3D orbital shaker at 4 C.
7. Put the tube in a magnetic holder, remove the supernatant, add 500 µLPBS with 0.1% SDS and 0.1% Triton-X100 and put in a 3D orbital shaker for 5 minutes.
8. Repeat the wash 1 more time.
9. Put the tube in a magnetic holder, remove the supernatant, add 500 µL PBS and put in a 3D orbital shaker for 5 minutes.
10. Repeat the wash 2 more times.
11. Resuspend with 100 µL of 100mM HEPES.

**Reduction:**

12. Add Dithiothreitol to a final concentration of 10mM
13. Incubate in a shaker for 60 min at 55C and 200 rpm.

**Alkylation:**

14. Immediately before use, add 100 mM HEPES to Iodoacetamide to a final concentration of 18.75mM Iodoacetamide.
15. Incubate for 30 min at room temperature covered with tinfoil**.**

**Trypsinization:**

16. Add 10nG mass spectrometry grade trypsin (2 µL of 5 nG / µL stock)
17. Incubate overnight at 37 C and 200rpm
18. After overnight digestion, stop trypsinization by adding 1 µL of Trifluoroacetic acid (TFA 0.1%)

**Sample multiplexing using TMT10plex™ Isobaric Label Reagent:**

19. Add 41 µL of the appropriate Peptide labelling reagent and incubate at room temperature for 1 hour.
20. Quench the labelling reaction with 8 µL of 5% Hydroxylamine at room temperature for 15 minutes.
21. Combine up to 10 samples in a single tube.
22. Leophelize the sample.
    a. Store the sample after lyophilization.
    b. Or resuspend with 100 mM of HEPES and continue to detergent removal.

**Solid Phase Extraction Stage Tips for detergent removal:**

**C18 Empore™ Stage - Tip:**

23. Use 3 layers of C18 Empore™ SPE disks,punched and inserted into 200 µL tips.
24. Activate with 100 µL MeOH
25. Centrifuge for 2 min at 1500 RCF and discard excess fluids.
26. Activate and cleaning from residual peptides with 100 µL buffer B (80% acetonitrile,0.1% TFA)
27. Centrifuge for 2 min at 1500 RCF and discard excess fluids.
28. Return to hydrophilic state with 100 µL buffer A (0.1%TFA)
29. Load up to 6 ug protein
30. Centrifuge for 2 min at 1500 RCF and discard excess fluids.
31. Return to hydrophilic state with 100 µL buffer A (0.1%TFA)
32. Repeat the previous step
33. Manually elute with 60 µL of buffer B.
34. Put the tube with the elution under vacuum.
35. Can be stored at room temperature.

**SCX strong cation exchange Empore™ Stage Tip:**

36. Use 3 layers of C18 Empore™ SPE disks.
37. Wash the layers with 400 µL of TFA 0.1%
38. Resuspend the sample with 100 µL TFA 0.1%
39. Load the sample and pass it through manually
40. Wash with 200 µL TFA 0.1%
41. Elution with 100 µL methanol 30% ammonium hydroxide 5%
42. Dry peptides with a speed-vac or a desiccator overnight

**Liquid Chromatography and mass spectrometry:**

43. Resolve peptides by reverse phase chromatography on 0.075 X 300 mm fused silica capillaries (J&W) packed with Reprosil reverse phase material (Dr Maisch GmbH, Germany).
44. Elute peptides with a 120 minutes linear gradient of 5% to 28% 15 minutes gradient of 28 to 95% and 25 minutes at 95% acetonitrile with 0.1% formic acid in water at flow rates of 0.15 µL/min.
45. Perform mass spectrometry by Q Exactive HF mass spectrometer (Thermo) in a positive mode (m/z 350–1600, resolution 120,000 for MS1 and 60,000 for MS2) using repetitively full MS scan followed by high collision dissociation (HCD, at 27 normalised collision energy) of the 10 most dominant ions (>1 charges) selected from the first MS scan.
46. The AGC settings were 3x10^6 for the full MS and 1x10^5 for the MS/MS scans. The intensity threshold for triggering MS/MS analysis was 8x10^4. A dynamic exclusion list was enabled with an exclusion duration of 20 seconds.

### Protocol - *S. mutans* Growth on extracted primary teeth:

#### Teeth preparation:

1. Incubate extracted primary molar in 5% Dakin's solution for 10 minutes.
2. Wash with sterilised double distilled water (DDW) for 10 minutes each.
3. Repeat 3 times.
4. Store in DDW at room temperature until use.

#### Saliva Preparation:

5. Spin down whole unstimulated saliva at 12,000 RCF for 7 minutes.
6. Keep the supernatant (Do not disturb the pellet).
7. Store in aliquots at -80 until use.

#### *Streptococcus Mutans* working solution:

8. Incubate Streptococcus mutans (S. mutans) in BHI until OD 0.6 - 0.8 is reached.
9. Prepare bacteria work solution: 5 uL of the prepared bacteria and 500 uL of Brain Heart Infusion (BHI) with 10uL of high / low DMFT saliva.
10. Place extracted prepared tooth in a 24 well plate, add 500 uL of the bacteria work solution.
11. Bacteria working solutions with 0.2% Chlorhexidine and PBS were used as negative and positive controls.
12. Incubate the sterilised extracted tooth in 500 uL of bacteria work solution for 5 minutes on an orbital shaker at room temperature.
13. Incubate for another 10 minutes with no shaking at room temperature.
14. Quick wash with 600 uL of sterilised DDW.
15. Repeat 3 times.
16. Incubate in 500 uL of BHI for 12 hours at 37C.
17. After incubation, wash twice with PBS and once with PBS 0.1% T.20 with 1:2,000 DAPI (of 5 mg/mL stock solution).
18. Analyse bacteria plaque formation under a microscope.

#### MS Sample preparations and analysis:

Briefly, Saliva antibodies from 34 participants were captured on protein L affinity beads. Solubilized *S. mutans* antigens (proteins) were allowed to bind the antibodies anchored to the beads. After washes, eluted proteins were digested, TMT labeled, multiplexed, cleaned and sent for identification by mass-spec analysis. From the identified proteins, we filtered out proteins with less than 3 peptides. To account for non-specific binding, a ratio of the relative intensity of the positive control (diluted protein lysate) to negative control (full protocol with antibodies omitted) was calculated.

**Supplementary 1:**

**Antigen Preparation:**
*Streptococcus Mutans* strain ATCC UA159 was obtained from Prof. Gilad Bachrach (The Hebrew University of Jerusalem, Israel). We confirmed it as S. mutans through targeted PCR using the primers TCGCGAAAAAGATAAACAAACA and GCCCCTTCACAGTTGGTTAG, sequencing of the resulting product, and protein mass-spectrometry. Bacteria were cultured according to ATCC guidelines (Streptococcus mutans Clarke). For antigen preparation, we cultivated the bacteria under anaerobic conditions in a Brain Heart Infusion (BHI) medium for 48 hours. Cells were centrifuged for 30 minutes at 2000 RFC using an Eppendorf 5910R centrifuge, medium discarded, cells were washed with PBS and pellet resuspended in RIPA lysis buffer, followed by sonication for 10 minutes at 20% power (OMNI International - Sonic Ruptor 400). Following centrifugation at 4500 RCF for 30 minutes, we discarded the pellet. Protein concentration was measured using nanodrop One$^{c}$ (Thermo-Fisher) 205 nm method. Proteins were aliquoted at 1 mG/mL and stored at -80°C until use.

**Supplementary 2:**

Plasmid construction expressing proteins of interest:

5 plasmids were engineered to express the following genes: Elongation factor Tu (P72483 Uniprot ID), UPF0246, GbpC (Q8DFT1 Uniprot ID), GbpB (Q8DWM3 Uniprot ID), Carbohydrate-binding domain-containing protein (Q8DWI5 Uniprot ID). The construction of the plasmids was based on the pDL278_P23-DsRed-Express2 vector's backbone (Shields et al. 2019). pDL278 vector was selected since it replicates and stays stable in various oral streptococci, besides being suitable for cloning and expression in *E. coli*. This vector incorporates the P23 promotor, allowing constitutive expression of genes, and a gene encoding for DsRed- Express2 protein, which is a red fluorescent protein. At the time of construction of plasmids of the concerned proteins, first their protein sequences were downloaded from uniprot, and the codon-optimized genes were cloned intopDL278_P23-DsRed-Express2 vector, replacing the DNA sequence of the gene expressing DsRed-Express2. To the C-terminal of the gene sequences, 6xHistag (HHHHHH peptide sequence) and Flag tag (DYKDDDDK peptide sequence) were added. HIS tag for easing purification and a flag tag for confident detection. Illustratration of the structure of plasmid expressing UPF0246 protein: The other plasmids of proteins of interest have similar structure but differ at the gene sequence region.

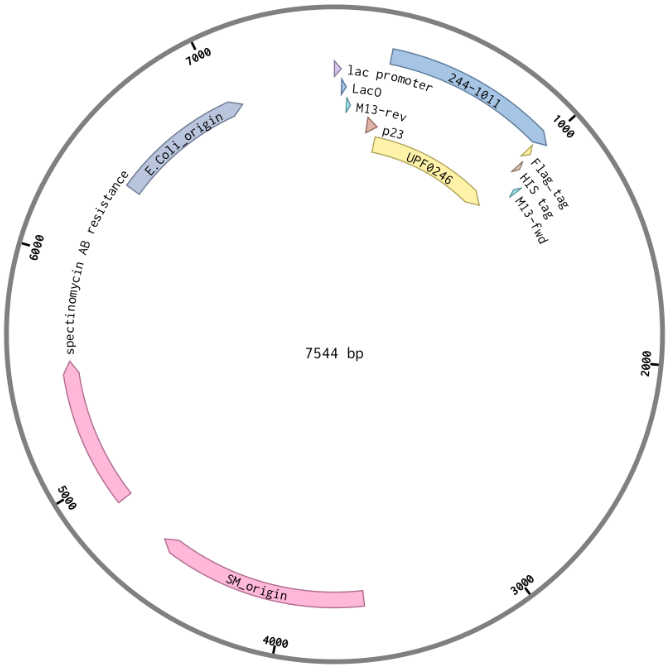


Bacterial protein expression:

The plasmids were transformed into *E. coli* strain BL21 for protein expression. One bacteria colony was selected from the petri dish and grown overnight in an LB medium with 50 µg/mL of spectinomycin at 37 °C and 180 x g speed. The following day, the expressed cells were collected and pelted by centrifugation at 4500 x g for 45 min at 4 °C, and pellets were frozen and stored at -80 °C.

Purification of proteins of *Streptococcus mutans* by Nickel beads:

The frozen pellets were lysed by suspending them with 20 mL lysis buffer made of 50mM phosphate buffer, 150 mM NaCl, 10 mM imidazole, one pill inhibitor protease cocktail (Mercury, REF. 11836170001), tween 20 0.5%, and 0.1 mM EDTA. After suspension, cells were disrupted by two sonication (OMNI International - Sonic Ruptor 400) cycles on ice, 8 minutes each (40 power level). The lysates were centrifuged at 4 °C for 30 min at max speed (Eppendorf 5910R, 20133 rcf). The supernatants, containing the proteins of interest, were collected and diluted with a 10 mL equilibration buffer and nickel (Ni) beads. The equilibration buffer is composed of 50 mM phosphate buffer, 150 mM NaCl and 10 mM imidazole. The supernatants along with the Ni beads were placed on a rotator at 4 °C for 4 hours of incubation, providing enough time for the Ni beads to bind to His- tagged proteins. Afterwards, they were centrifuged at 4 °C for 5 min at 500 x g. The supernatants were discarded and beads were resuspended in a 10 mL wash buffer, constituting 50 mM phosphate buffer, 150 mM NaCl, and 40 mM imidazole. The resuspensions were stirred at 4 °C for 15 minutes and later were centrifuged at 4 °C for 15 min at 500 x g. Again, the supernatants were aspirated and pellets were eluted by 5 mL elution buffer, made of equilibration buffer with 300 mM of imidazole. Then, the resuspensions were stirred at 4 °C for 15 minutes and finally were centrifuged at 4 °C for 15 min at 500 x g. The supernatants were collected and their protein concentration was measured with the aid of nanodrop OneC (Thermo-Fisher) by using the A280 method. The purification yields for elongation factor TU, UPF0246, glucan binding protein B and carbohydrate- binding domain- containing protein were 0.057 mg, 0.165 mg and 0.057 mg and 0.033 mg per 1 liter, respectively. By contrast, glucan binding protein C could not be purified either because E. coli BL21 strains failed to produce colonies, or failed to produce colonies which express the protein of interest. Trials with additional E. coli strains also failed. As a consequence, we concluded that the constitutive expression of this protein is toxic to E. coli.

A sample from the supernatants was taken in order to be analyzed via SDS gel and western blot to ensure the success of the process. The protein- containing solutions were diluted by 50% glycerol and then they were stored at -80 °C.

ELISA

The ELISA assay was performed using a compatible PVC clear high bind ELISA microtiter 96-well plate. First, coating was made with 100 µl per well of saliva samples collected from 2 groups of patients in ages between 6-12 years old; low DMFT (DMFT=0) patients and high DMFT (DMFT≥5). Prior to the binding step, the saliva samples were diluted 10 times by PBST

0.05%+ 1% BSA. The plate was placed on a rotator and incubated overnight at 4 °C. The next day, it was washed three times with 200 µl of PBS-T 0.05%. 200 µl Blocking solution containing PBST 0.05% and 5% BSA were added to each well, and the plate was incubated for 4 hours at RT on a rotator to prevent non-specific bindings. After incubation, the plate was inverted, the solution was flicked out, and washing steps were repeated. Binding was done with 100 µl of with the purified proteins, diluted to a desired concentration at a final volume of

100 µl per well. The plate was placed on a rotator and incubated for 40 minutes at RT. This was followed by two fast washes with PBST 0.05% and three long washes with PBST 0.05% + 1% BSA, each duration of wash was about 10 minutes waiting time on a rotator at RT.

Afterwards, 100 µl of anti- flag HRP conjugated antibody (1 µl antibody diluted into 5 mL PBST 0.05%) were pipetted to relevant wells and incubated for 30 minutes at RT. The last described washing steps were repeated and eventually 100 µL of TMB were added to each well and the plate was incubated at RT (4 – 30 min) for colour-reaction development. Finally, if colour develops, 50 µL of 0.5 M H2SO4 were added to each well to stop the enzymatic reaction. Plate absorbance was immediately measured using a plate reader at 450 nm absorbances. The previous describes a sandwich type of ELISA.

*S. mutans* tooth adherence assay:

The ability of saliva in preventing *S. mutans* attachment to teeth was measured by short incubation of cleaned teeth in BHI containing *S. mutans* in the presence of saliva from individuals with high (n = 10) or low (n = 10) DMFT scores according to this protocol 2. Teeth were washed and moved to fresh wells with growth medium but without bacteria, and adherent bacteria were allowed to grow overnight. The teeth were examined under LSM 900 microscope (Zeiss) using a 63X oil lens and 3-5 images were taken from each tooth. The images were scored by two blinded evaluators, asked to give a score of; 1 = no bacterial growth, 2 = low bacterial growth and 3 = high bacterial growth. The overall score of each tooth sample was calculated. Fisher's exact test was performed on the answers of the evaluators.

Bioinformatics analysis:

The 6 raw data files were searched against the Streptococcus Mutans UA159 proteome (uniprot #UP000002512) using MaxQuant V2.1.0.0 software (Max Planck institute). Protein minimal peptide count per protein was set to 3, with FDR set to 10% per peptide. Reverse sequences were used for decoy search and contaminant sequences were included in the search. Only proteins that included the relative intensity ratio of positive control / negative control > 1 were included in our data analysis.

#### Supplementary 3:

| file name | no bacteria | low density bacteria | high density bacteria | 1 = answe | correct? | 1 = yes | 0 = no | % correct | 68.32 | | |
|---|---|---|---|---|---|---|---|---|---|---|---|
| 1H_diff_Snap-3577 | | | 1 | | 1 | | | 100 | | 17 | high |
| 1H_Snap-3580 | | | 1 | | 1 | | | | | 17 | low |
| 1H_Snap-3581 | | | 1 | | 1 | | | | | | |
| 1H_Snap-3582 | | | 1 | | 1 | | | | | | |
| 1L_Snap-3583 | | 1 | | | 1 | | | 100 | | | |
| 1L_Snap-3584 | | 1 | | | 1 | | | | | | |
| 1L_Snap-3585 | | 1 | | | 1 | | | | | | |
| 2H_Snap-3586 | | 1 | | | 0 | | | 66 | | | |
| 2H_Snap-3587 | | | 1 | | 1 | | | | | | |
| 2H_Snap-3588 | | | 1 | | 1 | | | | | | |
| 2L_Snap-3589 | | 1 | | | 1 | | | 75 | | | |
| 2L_Snap-3590 | | | 1 | | 0 | | | | | | |
| 2L_Snap-3591 | | 1 | | | 1 | | | | | | |
| 2L_Snap-3592 | | 1 | | | 1 | | | | | | |
| 3H_Snap-3593 | | 1 | | | 0 | | | 50 | | | |
| 3H_Snap-3594 | | 1 | | | 0 | | | | | | |
| 3H_Snap-3595 | | | 1 | | 1 | | | | | | |
| 3H_Snap-3596 | | | 1 | | 1 | | | | | | |
| 3L_Snap-3597 | | 1 | | | 1 | | | 33.3 | | | |
| 3L_Snap-3598 | | | 1 | | 0 | | | | | | |
| 3L_Snap-3599 | | | 1 | | 0 | | | | | | |
| 4H_Snap-3600 | | | 1 | | 1 | | | 100 | | | |
| 4H_Snap-3601 | | | 1 | | 1 | | | | | | |
| 4H_Snap-3602 | | | 1 | | 1 | | | | | | |
| 4L_Snap-3 | 1 | | | | 1 | | | 100 | | | |
| 4L_Snap-3604 | | 1 | | | 1 | | | | | | |
| 4L_Snap-3605 | | 1 | | | 1 | | | | | | |
| 4L_Snap-3606 | | 1 | | | 1 | | | | | | |
| 5H_Snap-3607 | | | 1 | | 1 | | | 100 | | | |
| 5H_Snap-3608 | | | 1 | | 1 | | | | | | |
| 5H_Snap-3609 | | | 1 | | 1 | | | | | | |
| 5L_Snap-3610 | | | 1 | | 0 | | | 33.3 | | | |
| 5L_Snap-3611 | | | 1 | | 0 | | | | | | |
| 5L_Snap-3612 | | 1 | | | 1 | | | | | | |

| high=1 low =0 | file name | no bacteria | low density bacteria | high density bacteria | 1 = answer | correct? | 1 = yes | 0 = no | % correct |
|---|---|---|---|---|---|---|---|---|---|
| 1 | 1H_diff_Snap-3577 | | | 1 | | 1 | | | 100 |
| 1 | 1H_Snap-3580 | | | 1 | | 1 | | | |
| 1 | 1H_Snap-3581 | | | 1 | | 1 | | | |
| 1 | 1H_Snap-3582 | | | 1 | | 1 | | | |
| 0 | 1L_Snap-3583 | | 1 | | | 1 | | | 100 |
| 0 | 1L_Snap-3584 | | 1 | | | 1 | | | |
| 0 | 1L_Snap-3585 | | 1 | | | 1 | | | |
| 1 | 2H_Snap-3586 | | 1 | | | 0 | | | 66.6 |
| 1 | 2H_Snap-3587 | | | 1 | | 1 | | | |
| 1 | 2H_Snap-3588 | | | 1 | | 1 | | | |
| 0 | 2L_Snap-3589 | | 1 | | | 1 | | | 75 |
| 0 | 2L_Snap-3590 | | | 1 | | 0 | | | |
| 0 | 2L_Snap-3591 | | 1 | | | 1 | | | |
| 0 | 2L_Snap-3592 | | 1 | | | 1 | | | |
| 1 | 3H_Snap-3593 | | 1 | | | 0 | | | 100 |
| 1 | 3H_Snap-3594 | | 1 | | | 0 | | | |
| 1 | 3H_Snap-3595 | | | 1 | | 1 | | | |
| 1 | 3H_Snap-3596 | | | 1 | | 1 | | | |
| 0 | 3L_Snap-3597 | | 1 | | | 1 | | | 66.6 |
| 0 | 3L_Snap-3598 | | 1 | | | 1 | | | |
| 0 | 3L_Snap-3599 | | | 1 | | 0 | | | |
| 1 | 4H_Snap-3600 | | | 1 | | 1 | | | 100 |
| 1 | 4H_Snap-3601 | | | 1 | | 1 | | | |
| 1 | 4H_Snap-3602 | | | 1 | | 1 | | | |
| 0 | 4L_Snap-3603 | 1 | | | | 1 | | | 75 |
| 0 | 4L_Snap-3604 | | | 1 | | 0 | | | |
| 0 | 4L_Snap-3605 | | 1 | | | 1 | | | |
| 0 | 4L_Snap-3606 | | 1 | | | 1 | | | |
| 1 | 5H_Snap-3607 | | | 1 | | 1 | | | 100 |
| 1 | 5H_Snap-3608 | | | 1 | | 1 | | | |
| 1 | 5H_Snap-3609 | | | 1 | | 1 | | | |
| 0 | 5L_Snap-3610 | | | 1 | | 0 | | | 33.3 |
| 0 | 5L_Snap-3611 | | | 1 | | 0 | | | |
| 0 | 5L_Snap-3612 | | 1 | | | 1 | | | |

| Results | | | |
|---|---|---|---|
| | LOW | HIGH | ***Marginal Row Totals*** |
| GUESS HIGH | 5 | 14 | 19 |
| GUESS LOW | 12 | 3 | 15 |
| ***Marginal Column Totals*** | 17 | 17 | 34 (Grand Total) |

| | Caries free | High DMFT |
|---|---|---|
| High DMFT evaluated | 5 | 14 |
| Caries free evaluated | 12 | 3 |

P.value = 0.0049

* both evaluators

**Supplementary 4:**
**Interactions map:**

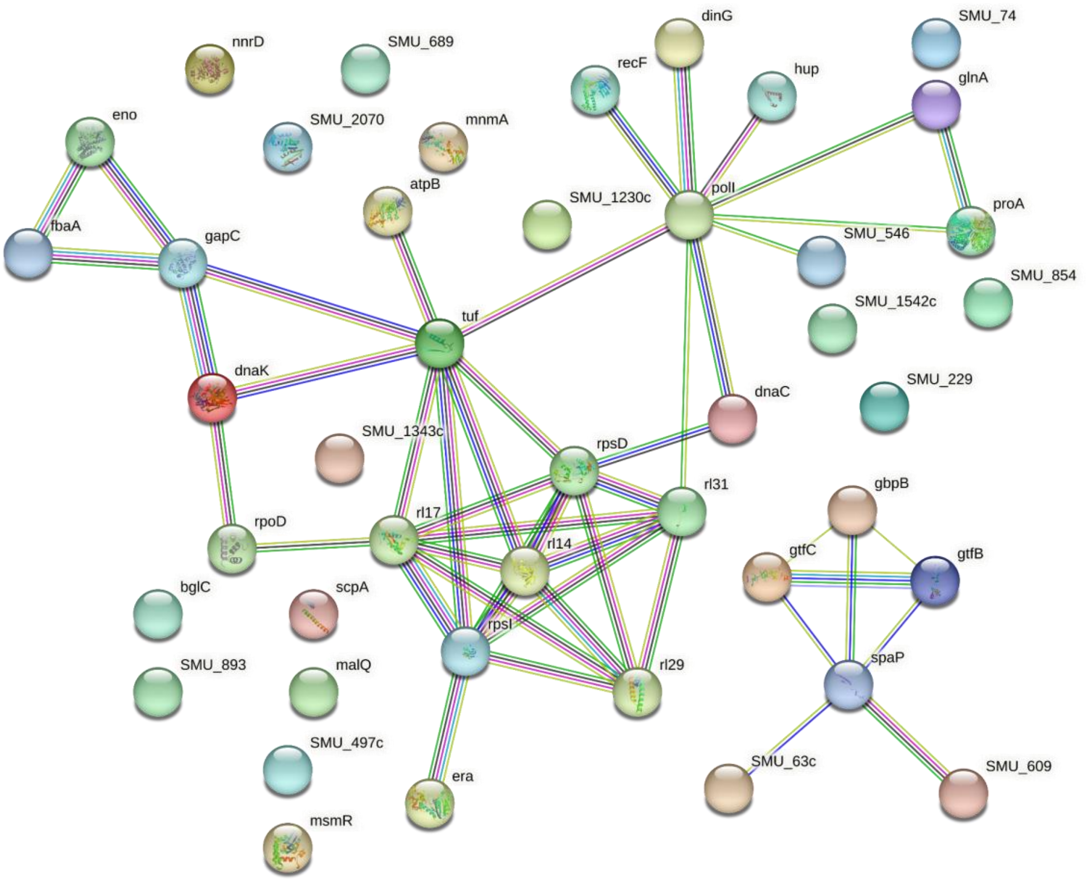


**String protein annotations:**

#node identifier domain_summary_url annotation other_names_and_aliases
SMU_1230c 210007.SMU_1230c Conserved hypothetical protein; Uncharacterized protein; Best Blastp Hit: dbj|BAA94663.1| (D86934) ORF N043 [Staphylococcus aureus] https://smart.embl.de/smart/DDvec.cgi?smart=158:Pfam_DUF1643(18|146)+ 1028526,24377608,AAN58915,AAN58915.1,Conserved hypothetical protein,GI:24377608,Hypothetical protein,NC_004350.2,NP_721609,NP_721609.1,Q8DTU3,Q8DTU3_STRMU,SMU_1230c,Uncharacterized protein,WP_002263212.1,conserved hypothetical protein,smu:SMU_1230c
SMU_1343c 210007.SMU_1343c Putative polyketide synthase; Best Blastp Hit: gb|AAF08795.1|AF184956_2 (AF184956) MycA [Bacillus subtilis] https://smart.embl.de/smart/DDvec.cgi?smart=1104:Pfam_ketoacyl-synt(1|238)+Pfam_Ketoacyl-synt_C(246|364)+Pfam_KAsynt_C_assoc(366|484)+Pfam_PP-binding(591|654)+Pfam_Condensation(675|1099)+ 1028611,24377719,AAN59016,AAN59016.1,GI:24377719,NC_004350.2,NP_721710,NP_721710.1,Polyketide synthase,Putative polyketide synthase,Q8DTJ4,Q8DTJ4_STRMU,SMU_1343c,WP_011074616.1,putative polyketide synthase,smu:SMU_1343c
SMU_1542c 210007.SMU_1542c Uncharacterized protein; Best Blastp Hit: pir||F69795 conserved hypothetical protein yerQ - Bacillus subtilis >gi|2632986|emb|CAB12492.1| (Z99107) similar to hypothetical proteins [Bacillus subtilis]

https://smart.embl.de/smart/DDvec.cgi?smart=347:DAGKc(7|138)+
1028785,2.7.1.107,24377911,AAN59191,AAN59191.1,Conserved hypothetical protein,DAGKc domain-containing protein,Diacylglycerol kinase (ATP) ,EC 2.7.1.107,GI:24377911,Lipid kinase,NC_004350.2,NP_721885,NP_721885.1,Q8DT50,Q8DT50_STRMU,SMU_1542c,WP_002262954.1,conserved hypothetical protein,smu:SMU_1542c
SMU_2070	210007.SMU_2070	Conserved hypothetical protein; Belongs to the UPF0246 family
https://smart.embl.de/smart/DDvec.cgi?smart=242:Pfam_H2O2_YaaD(1|229)+
1029247,24378429,AAN59666,AAN59666.1,Conserved hypothetical protein,GI:24378429,Hypothetical protein,NC_004350.2,NP_722360,NP_722360.1,Q8DRY6,SMU_2070,UPF0246 protein SMU_2070,Uncharacterized protein,WP_002262377.1,Y2070_STRMU,conserved hypothetical protein,smu:SMU_2070
SMU_229	210007.SMU_229	Uncharacterized protein; Best Blastp Hit: pir||E69879 conserved hypothetical protein yloV - Bacillus subtilis >gi|2337813|emb|CAA74257.1| (Y13937) YloV protein [Bacillus subtilis] >gi|2633956|emb|CAB13457.1| (Z99112) similar to hypothetical proteins [Bacillus subtilis]
https://smart.embl.de/smart/DDvec.cgi?smart=555:Dak2(34|199)+Dak1_2(240|555)+
1029500,24376607,AAN58001,AAN58001.1,Conserved hypothetical protein,DhaL domain-containing protein,GI:24376607,Hypothetical protein,NC_004350.2,NP_720695,NP_720695.1,Q8DW46,Q8DW46_STRMU,SMU_229,Uncharacterized protein,WP_002262763.1,conserved hypothetical protein,smu:SMU_229
SMU_497c	210007.SMU_497c	Uncharacterized protein; Best Blastp Hit: sp|P32437|YVYE_BACSU HYPOTHETICAL 24.8 KD PROTEIN IN DEGS-TAGO INTERGENIC REGION >gi|7429381|pir||A70049 conserved hypothetical protein yvyE - Bacillus subtilis >gi|1762328|gb|AAC44936.1| (U56901) Ycr59c/YigZ homolog [Bacillus subtilis] >gi|2636077|emb|CAB15568.1| (Z99122) alternate gene name: yvhK similar to hypothetical proteins [Bacillus subtilis]
https://smart.embl.de/smart/DDvec.cgi?smart=209:Pfam_UPF0029(17|122)+Pfam_DUF1949(138|193)+	1029589,24376871,AAN58242,AAN58242.1,Conserved hypothetical protein,GI:24376871,Hypothetical protein,NC_004350.2,NP_720936,NP_720936.1,Q8DVJ1,Q8DVJ1_STRMU,SMU_497c,Uncharacterized protein,WP_002263027.1,conserved hypothetical protein,smu:SMU_497c
SMU_546	210007.SMU_546	Putative GTP-binding protein; Best Blastp Hit: dbj|BAB06351.1| (AP001516) GTP-binding protein TypA/BipA (tyrosine phosphorylated protein A) [Bacillus halodurans]
https://smart.embl.de/smart/DDvec.cgi?smart=614:Pfam_GTP_EFTU(6|201)+Pfam_GTP_EFTU_D2(224|294)+EFG_C(403|485)+
1028019,24376921,AAN58288,AAN58288.1,GI:24376921,GTP-binding protein,NC_004350.2,NP_720982,NP_720982.1,Putative GTP-binding protein,Q8DVE5,Q8DVE5_STRMU,SMU_546,WP_002262063.1,putative GTP-binding protein,smu:SMU_546
SMU_609	210007.SMU_609	Best Blastp Hit: pir||A60328 40K cell wall protein precursor (sr 5' region) - Streptococcus mutans (strain OMZ175, serotype f)
https://smart.embl.de/smart/DDvec.cgi?smart=611:Pfam_YSIRK_signal(9|33)+SH3b(187|253)+SH3b(267|333)+Pfam_GBS_Bsp-like(343|434)+Pfam_GBS_Bsp-like(447|538)+SH3b(550|608)+	1029570,24376985,40K cell wall

protein,AAN58347,AAN58347.1,GI:24376985,NC_004350.2,NP_721041,NP_721041.1,Putative 40K cell wall protein,Putative 40K cell wall protein precursor,Q8CVC7,Q8CVC7_STRMU,SMU_609,WP_002352220.1,putative 40K cell wall protein precursor,smu:SMU_609
SMU_63c	210007.SMU_63c	Conserved hypothetical protein; Uncharacterized protein; Best Blastp Hit: pir||T33369 hypothetical protein H02F09.3 - Caenorhabditis elegans >gi|3319425|gb|AAC64622.1| (AF077538) unknown [Caenorhabditis elegans]
	https://smart.embl.de/smart/DDvec.cgi?smart=613:Pfam_Cthe_2159(384|471)+
	1029648,24376442,AAN57850,AAN57850.1,Conserved hypothetical protein,GI:24376442,Hypothetical protein,NC_004350.2,NP_720544,NP_720544.1,Q8DWI5,Q8DWI5_STRMU,SMU_63c,Uncharacterized protein,WP_002263401.1,conserved hypothetical protein,smu:SMU_63c
SMU_689	210007.SMU_689	Lysozyme; Best Blastp Hit: pir||C60328 hypothetical protein 2 (sr 5' region) - Streptococcus mutans (strain OMZ175, serotype f)
	https://smart.embl.de/smart/DDvec.cgi?smart=979:SIGNAL(1|24)+Pfam_GBS_Bsp-like(183|266)+Pfam_GBS_Bsp-like(274|363)+Pfam_GBS_Bsp-like(377|467)+Pfam_GBS_Bsp-like(483|560)+Pfam_GBS_Bsp-like(568|657)+Pfam_GBS_Bsp-like(670|760)+Pfam_Glyco_hydro_25(776|964)+
	1028106,24377068,3.2.1.17,AAN58422,AAN58422.1,GI:24377068,Hypothetical protein,Lysozyme,NC_004350.2,NP_721116,NP_721116.1,Q8DV28,Q8DV28_STRMU,SMU_689,WP_002263335.1,hypothetical protein,smu:SMU_689
SMU_74	210007.SMU_74	Conserved hypothetical protein; Best Blastp Hit: pir||S56619 gpmB protein - Escherichia coli >gi|537235|gb|AAA97291.1| (U14003) Kenn Rudd identifies as gpmB [Escherichia coli] >gi|1790856|gb|AAC77348.1| (AE000509) phosphoglyceromutase 2 [Escherichia coli]
	https://smart.embl.de/smart/DDvec.cgi?smart=236:PGAM(5|182)+
	1029660,24376453,AAN57860,AAN57860.1,Conserved hypothetical protein,GI:24376453,Hypothetical protein,NC_004350.2,NP_720554,NP_720554.1,Q8DWH6,Q8DWH6_STRMU,SMU_74,Uncharacterized protein,WP_002263410.1,conserved hypothetical protein,smu:SMU_74
SMU_854	210007.SMU_854	23S rRNA pseudouridine1911/1915/1917 synthase; Responsible for synthesis of pseudouridine from uracil
	https://smart.embl.de/smart/DDvec.cgi?smart=296:S4(10|73)+Pfam_PseudoU_synth_2(82|236)+	1029411,23S rRNA pseudouridine1911/1915/1917 synthase ,24377230,5.4.99.-,5.4.99.23,AAN58570,AAN58570.1,EC 5.4.99.23,GI:24377230,NC_004350.2,NP_721264,NP_721264.1,Pseudouridine synthase,Pseudouridylate synthase,Putative pseudouridylate synthase,Q8DUP9,Q8DUP9_STRMU,SMU_854,WP_002262003.1,putative pseudouridylate synthase,smu:SMU_854
SMU_893	210007.SMU_893	Putative anticodon nuclease; Best Blastp Hit: pir||H81152 anticodon nuclease NMB0832 [imported] - Neisseria meningitidis (group B strain MD58) >gi|7226066|gb|AAF41243.1| (AE002436) anticodon nuclease [Neisseria meningitidis MC58] https://smart.embl.de/smart/DDvec.cgi?smart=381:Pfam_AAA_13(176|373)+
	1028259,24377271,AAN58607,AAN58607.1,Anticodon nuclease,GI:24377271,NC_004350.2,NP_721301,NP_721301.1,Putative anticodon nuclease,Q8DUM5,Q8DUM5_STRMU,SMU_893,WP_002262887.1,putative anticodon nuclease,smu:SMU_893

atpB 210007.SMU_1528 FoF1 membrane-bound proton-translocating ATPase, beta subunit; Produces ATP from ADP in the presence of a proton gradient across the membrane. The catalytic sites are hosted primarily by the beta subunits
https://smart.embl.de/smart/DDvec.cgi?smart=468:Pfam_ATP-synt_ab_N(6|77)+AAA(147|332)+
1028766,24377897,7.1.2.2,7.2.2.1,AAN59178,AAN59178.1,ATP synthase F0F1 subunit beta,ATP synthase F1 sector subunit beta,ATP synthase subunit beta,ATPB_STRMU,AtpB,EC 7.1.2.2,EC 7.2.2.1,F-ATPase subunit beta,F-type H+/Na+-transporting ATPase subunit beta ,GI:24377897,NC_004350.2,NP_721872,NP_721872.1,P95789,SMU_1528,WP_002262940.1,atpB,atpD,smu:SMU_1528
bglC 210007.SMU_983 Putative transcriptional regulator; Best Blastp Hit: gb|AAF89977.1|AF206272_3 (AF206272) transcriptional regulator [Streptococcus mutans]
https://smart.embl.de/smart/DDvec.cgi?smart=301:Pfam_AraC_binding(27|172)+HTH_ARAC(210|293)+
1028315,24377355,AAN58684,AAN58684.1,BglC,GI:24377355,NC_004350.2,NP_721378,NP_721378.1,Putative transcriptional regulator,Q8DUE9,Q8DUE9_STRMU,SMU_983,Transcriptional regulator,WP_011074572.1,bglC,putative transcriptional regulator,smu:SMU_983
dinG 210007.SMU_1313c Bifunctional atp-dependent dna helicase/dna polymerase iii subunit epsilon; 3'-5' exonuclease
https://smart.embl.de/smart/DDvec.cgi?smart=820:EXOIII(8|172)+DEXDc(242|506)+HELICc(650|786)+ 1028587,24377689,3'-5' exonuclease DinG,3.1.-.-,3.6.4.12,AAN58990,AAN58990.1,ATP-dependent DNA helicase DinG ,EC 3.6.4.12,GI:24377689,NC_004350.2,NP_721684,NP_721684.1,Putative ATP-dependent DNA helicase,Q8DTM0,Q8DTM0_STRMU,SMU_1313c,WP_011074609.1,dinG,putative ATP-dependent DNA helicase,smu:SMU_1313c
dnaC 210007.SMU_2138 Putative replicative dna helicase (dna polymerase iii delta prime subunit); Participates in initiation and elongation during chromosome replication; it exhibits DNA-dependent ATPase activity
https://smart.embl.de/smart/DDvec.cgi?smart=454:Pfam_DnaB(9|111)+Pfam_AAA_25(171|368)+ 1029303,24378497,3.6.4.12,AAN59729,AAN59729.1,DnaC,EC 3.6.4.12,GI:24378497,NC_004350.2,NP_722423,NP_722423.1,Q8DRS9,Q8DRS9_STRMU,Replicative DNA helicase,Replicative DNA helicase ,SMU_2138,WP_002262440.1,dnaC,smu:SMU_2138
dnaK 210007.SMU_82 Heat shock protein, dnak (hsp-70); Acts as a chaperone
https://smart.embl.de/smart/DDvec.cgi?smart=612:Pfam_MreB_Mbl(2|352)+
1029666,24376461,AAN57867,AAN57867.1,Chaperone protein DnaK,DNAK_STRMU,DnaK,GI:24376461,HSP70,Heat shock 70 kDa protein,Heat shock protein 70,Heat shock protein, DnaK (HSP-70),Molecular chaperone DnaK,NC_004350.2,NP_720561,NP_720561.1,O06942,SMU_82,WP_002263417.1,dnaK,heat shock protein, DnaK (HSP-70),smu:SMU_82
eno 210007.SMU_1247 Putative enolase; Catalyzes the reversible conversion of 2-phosphoglycerate into phosphoenolpyruvate. It is essential for the degradation of carbohydrates via glycolysis (By similarity). Binds plasminogen and human salivary mucin MG2 when expressed on the bacterial cell surface, potentially allowing the bacterium to acquire surface-associated proteolytic activity that may help the dissemination through oral tissues and entrance into the blood stream

https://smart.embl.de/smart/DDvec.cgi?smart=432:Enolase_N(4|134)+Enolase_C(139|429)+ 1028540,2-phospho-D-glycerate hydro-lyase,2-phosphoglycerate dehydratase,24377625,4.2.1.11,AAN58930,AAN58930.1,EC 4.2.1.11,ENO_STRMU,Eno,Enolase,Enolase ,GI:24377625,NC_004350.2,NP_721624,NP_721624.1,Putative enolase,Q8DTS9,SMU_1247,WP_002263196.1,eno,putative enolase,smu:SMU_1247
era 210007.SMU_1617 Gtp-binding protein; An essential GTPase that binds both GDP and GTP, with rapid nucleotide exchange. Plays a role in 16S rRNA processing and 30S ribosomal subunit biogenesis and possibly also in cell cycle regulation and energy metabolism (By similarity). Has GTPase activity, binds both GDP and GTP, does not bind UTP, CTP or ATP
https://smart.embl.de/smart/DDvec.cgi?smart=299:Pfam_GTP_EFTU(6|171)+Pfam_KH_2(205|283)+
1028847,24377985,AAN59258,AAN59258.1,ERA_STRMU,Era,GI:24377985,GTP-binding protein,GTPase,GTPase Era,NC_004350.2,NP_721952,NP_721952.1,P37214,SGP,SMU_1617,WP_002262773.1,era,smu:SMU_1617,spg
fbaA 210007.SMU_99 Fructose-bisphosphate aldolase, class ii; Best Blastp Hit: dbj|BAB16889.1| (AB050113) class-II aldolase [Streptococcus bovis]
https://smart.embl.de/smart/DDvec.cgi?smart=293:Pfam_F_bP_aldolase(4|292)+
1029679,24376476,4.1.2.13,AAN57881,AAN57881.1,EC 4.1.2.13,FbaA,Fructose-1,6-biphosphate aldolase,Fructose-bisphosphate aldolase,Fructose-bisphosphate aldolase, class II ,GI:24376476,NC_004350.2,NP_720575,NP_720575.1,Q8DWG0,Q8DWG0_STRMU,SMU_99,WP_002263577.1,fbaA,fructose-1,6-biphosphate aldolase,smu:SMU_99
gapC 210007.SMU_360 Extracellular glyceraldehyde-3-phosphate dehydrogenase; Belongs to the glyceraldehyde-3-phosphate dehydrogenase family
https://smart.embl.de/smart/DDvec.cgi?smart=337:Gp_dh_N(3|153)+Pfam_Gp_dh_C(158|315)+ 1.2.1.-,1.2.1.12,1027909,24376735,AAN58118,AAN58118.1,EC 1.2.1.12,GI:24376735,GapC,Glyceraldehyde 3-phosphate dehydrogenase ,Glyceraldehyde-3-phosphate dehydrogenase,NC_004350.2,NP_720812,NP_720812.1,Q8DVV3,Q8DVV3_STRMU,SMU_360,WP_002262489.1,gapC,smu:SMU_360
gbpB 210007.SMU_22 Peptidoglycan dl-endopeptidase cwlo; Putative secreted antigen GbpB/SagA putative peptidoglycan hydrolase; Best Blastp Hit: gb|AAD00288.1| (U78607) putative secreted protein [Streptococcus mutans]
https://smart.embl.de/smart/DDvec.cgi?smart=431:SIGNAL(1|19)+COIL(32|99)+COIL(144|268)+Pfam_CHAP(327|406)+ 1029610,24376400,3.4.-.-,AAN57811,AAN57811.1,EC 3.4.-.-,GI:24376400,GbpB,NC_004350.2,NP_720505,NP_720505.1,Peptidoglycan DL-endopeptidase CwlO ,Putative secreted antigen GbpB/SagA,Q8DWM3,Q8DWM3_STRMU,SMU_22,Secreted antigen GbpB/SagA,WP_002263140.1,gbpB,putative secreted antigen GbpB/SagA,sagA,smu:SMU_22
glnA 210007.SMU_364 Glutamine synthetase type 1; Belongs to the glutamine synthetase family https://smart.embl.de/smart/DDvec.cgi?smart=448:Pfam_Gln-synt_N(16|100)+Gln-synt_C(106|358)+
1029365,24376739,6.3.1.2,AAN58122,AAN58122.1,EC 6.3.1.2,GI:24376739,GlnA,Glutamate--ammonia ligase,Glutamine synthetase ,Glutamine

synthetase type 1,NC_004350.2,NP_720816,NP_720816.1,Q8DVU9,Q8DVU9_STRMU,SMU_364,WP_002262485.1,glnA,glutamine synthetase type 1,smu:SMU_364
gtfB 210007.SMU_1004 Glucosyltransferase-i; Production of extracellular glucans, that are thought to play a key role in the development of the dental plaque because of their ability to adhere to smooth surfaces and mediate the aggregation of bacterial cells and food debris
https://smart.embl.de/smart/DDvec.cgi?smart=1476:Pfam_Glyco_hydro_70(269|1074)+Pfam_CW_binding_1(1088|1106)+Pfam_CW_binding_1(1280|1299)+Pfam_CW_binding_1(1345|1364)+Pfam_CW_binding_1(1410|1429)+
1028336,2.4.1.5,24377379,AAN58705,AAN58705.1,Dextransucrase,Dextransucrase,EC 2.4.1.5,GI:24377379,GTF-I,GTFB_STRMU,Glucosyltransferase-I,GtfB,NC_004350.2,NP_721399,NP_721399.1,O69381,O69384,O69387,O69390,O69396,P08987,SMU_1004,Sucrose 6-glucosyltransferase,WP_002352268.1,glucosyltransferase-I,gtfB,smu:SMU_1004
gtfC 210007.SMU_1005 Glucosyltransferase-si; Production of extracellular glucans, that are thought to play a key role in the development of the dental plaque because of their ability to adhere to smooth surfaces and mediate the aggregation of bacterial cells and food debris
https://smart.embl.de/smart/DDvec.cgi?smart=1455:Pfam_Glyco_hydro_70(295|1103)+Pfam_CW_binding_1(1117|1135)+Pfam_CW_binding_1(1373|1392)+
1028343,2.4.1.5,24377380,3AIB,3AIC,3AIE,AAN58706,AAN58706.1,Dextransucrase,Dextransucrase ,EC 2.4.1.5,GI:24377380,GTF-SI,GTFC_STRMU,Glucosyltransferase-SI,GtfC,NC_004350.2,NP_721400,NP_721400.1,O69382,O69385,O69388,O69391,O69397,P05427,P13470,SMU_1005,Sucrose 6-glucosyltransferase,WP_002352269.1,glucosyltransferase-SI,gtfC,smu:SMU_1005
hup 210007.SMU_589 Putative dna-binding protein; Histone-like DNA-binding protein which is capable of wrapping DNA to stabilize it, and thus to prevent its denaturation under extreme environmental conditions. Seems also to act as a fortuitous virulence factor in delayed sequelae by binding to heparan sulfate-proteoglycans in the extracellular matrix of target organs and acting as a nidus for in situ immune complex formation
https://smart.embl.de/smart/DDvec.cgi?smart=91:BHL(2|91)+
1028043,24376965,5FBM,AAN58328,AAN58328.1,DBH_STRMU,DNA-binding protein,DNA-binding protein HU,DNA-binding protein HU-beta,GI:24376965,NC_004350.2,NP_721022,NP_721022.1,Putative DNA-binding protein,Q9XB21,SMU_589,WP_002262099.1,hlpA,hup,putative DNA-binding protein,smu:SMU_589
malQ 210007.SMU_1565 Putative 4-alpha-glucanotransferase; 4-alpha-glucanotransferase; Best Blastp Hit: sp|P29851|MALQ_STRPN 4-ALPHA-GLUCANOTRANSFERASE (AMYLOMALTASE) (DISPROPORTIONATING ENZYME) (D-ENZYME) >gi|153704|gb|AAA26923.1| (J01796) amylomaltase [Streptococcus pneumoniae]
https://smart.embl.de/smart/DDvec.cgi?smart=509:Pfam_Glyco_hydro_77(10|480)+
1028801,2.4.1.25,24377932,4-alpha-glucanotransferase,4-alpha-glucanotransferase,AAN59211,AAN59211.1,Amylomaltase,Disproportionating enzyme,EC 2.4.1.25,GI:24377932,MalQ,NC_004350.2,NP_721905,NP_721905.1,Putative 4-alpha-glucanotransferase,Q8DT30,Q8DT30_STRMU,SMU_1565,WP_002262974.1,malQ,putative 4-alpha-glucanotransferase,smu:SMU_1565
mnmA 210007.SMU_2143c Trna-specific 2-thiouridylase mnma; Catalyzes the 2-thiolation of uridine at the wobble position (U34) of tRNA, leading to the formation of s(2)U34

https://smart.embl.de/smart/DDvec.cgi?smart=373:Pfam_NAD_synthase(1|128)+ 1029315,2.8.1.13,24378503,AAN59734,AAN59734.1,EC 2.8.1.13,GI:24378503,MNMA_STRMU,NC_004350.2,NP_722428,NP_722428.1,Putative tRNA,Q8DRS4,SMU_2143c,TRNA-specific 2-thiouridylase MnmA,TRNA-uridine 2-sulfurtransferase ,WP_002262445.1,mnmA,putative tRNA,smu:SMU_2143c,tRNA-specific 2-thiouridylase MnmA,trmU

msmR 210007.SMU_876 Regulatory protein for the msm operon for multiple sugar metabolism. Activates the transcription of the msmEFGK, aga, dexB and gftA genes https://smart.embl.de/smart/DDvec.cgi?smart=278:Pfam_AraC_binding(18|152)+HTH_ARAC(189|272)+ 1028226,24377254,AAN58591,AAN58591.1,GI:24377254,MSM operon regulatory protein,MSMR_STRMU,Msm operon regulatory protein,MsmR,NC_004350.2,NP_721285,NP_721285.1,Putative MSM operon regulatory protein,Q00753,SMU_876,WP_002262870.1,msmR,putative MSM operon regulatory protein,smu:SMU_876

nnrD 210007.SMU_573 Adp-dependent nad(p)h-hydrate dehydratase; Catalyzes the dehydration of the S-form of NAD(P)HX at the expense of ADP, which is converted to AMP. Together with NAD(P)HX epimerase, which catalyzes the epimerization of the S-and R-forms, the enzyme allows the repair of both epimers of NAD(P)HX, a damaged form of NAD(P)H that is a result of enzymatic or heat-dependent hydration https://smart.embl.de/smart/DDvec.cgi?smart=277:Pfam_Carb_kinase(27|270)+ 1029441,24376950,3BGK,4.2.1.136,AAN58314,AAN58314.1,ADP-dependent (S)-NAD(P)H-hydrate dehydratase,ADP-dependent NAD(P)H-hydrate dehydratase ,ADP-dependent NAD(P)HX dehydratase,Conserved hypothetical protein,EC 4.2.1.136,GI:24376950,Hypothetical protein,NC_004350.2,NP_721008,NP_721008.1,Q8DVC0,Q8DVC0_STRMU,SMU_573,WP_002263560.1,conserved hypothetical protein,nnrD,smu:SMU_573

polI 210007.SMU_297 Dna polymerase i (pol i); In addition to polymerase activity, this DNA polymerase exhibits 5'-3' exonuclease activity https://smart.embl.de/smart/DDvec.cgi?smart=878:HhH2(180|215)+35EXOc(288|466)+POLAc(634|842)+ 1029369,2.7.7.7,24376673,AAN58061,AAN58061.1,DNA polymerase I,DNA polymerase I ,DNA polymerase I (POL I),EC 2.7.7.7,GI:24376673,NC_004350.2,NP_720755,NP_720755.1,PolI,Q8DVZ5,Q8DVZ5_STRMU,SMU_297,WP_002263533.1,polA,polI,smu:SMU_297

proA 210007.SMU_450 Putative gamma-glutamyl phosphate reductase; Catalyzes the NADPH-dependent reduction of L-glutamate 5- phosphate into L-glutamate 5-semialdehyde and phosphate. The product spontaneously undergoes cyclization to form 1-pyrroline-5-carboxylate https://smart.embl.de/smart/DDvec.cgi?smart=416:Pfam_Aldedh(4|296)+Pfam_Aldedh(294|406)+ 1.2.1.41,1029582,24376825,AAN58200,AAN58200.1,EC 1.2.1.41,GI:24376825,GPR,GSA dehydrogenase,Gamma-glutamyl phosphate reductase,Glutamate-5-semialdehyde dehydrogenase,Glutamate-5-semialdehyde dehydrogenase ,Glutamyl-gamma-semialdehyde dehydrogenase,NC_004350.2,NP_720894,NP_720894.1,PROA_STRMU,ProA,Putative gamma-glutamyl phosphate reductase,Q8DVM9,SMU_450,WP_002262086.1,proA,putative gamma-glutamyl phosphate reductase,smu:SMU_450

recF 210007.SMU_2156 Putative recf protein, atpase involved in dna repair; The RecF protein is involved in DNA metabolism; it is required for DNA replication and normal SOS inducibility. RecF binds preferentially to single-stranded, linear DNA. It also seems to bind

ATP https://smart.embl.de/smart/DDvec.cgi?smart=363:Pfam_AAA_15(1|337)+ 1029328,24378515,AAN59745,AAN59745.1,DNA replication and repair protein RecF,GI:24378515,NC_004350.2,NP_722439,NP_722439.1,Q8DRR3,RECF_STRMU,RecF,Recombination protein F,SMU_2156,WP_002262456.1,recF,smu:SMU_2156

rl14 210007.SMU_2017 Large subunit ribosomal protein l14; Binds to 23S rRNA. Forms part of two intersubunit bridges in the 70S ribosome
https://smart.embl.de/smart/DDvec.cgi?smart=122:Ribosomal_L14(1|122)+
1029205,24378378,50S ribosomal protein L14,AAN59620,AAN59620.1,GI:24378378,Large subunit ribosomal protein L14,NC_004350.2,NP_722314,NP_722314.1,Q8DS23,RL14_STRMU,Rl14,SMU_2017,WP_002262330.1,rl14,rplN,smu:SMU_2017

rl17 210007.SMU_2000 Large subunit ribosomal protein l17; Best Blastp Hit: sp|P20277|RL17_BACSU 50S RIBOSOMAL PROTEIN L17 (BL15) (BL21) >gi|80370|pir||F32307 ribosomal protein L17 - Bacillus subtilis >gi|142464|gb|AAA22218.1| (M26414) ribosomal protein L17 [Bacillus subtilis] >gi|1044992|gb|AAB06827.1| (L47971) ribosomal protein L17 [Bacillus subtilis] >gi|2632411|emb|CAB11920.1| (Z99104) ribosomal protein L17 (BL15) [Bacillus subtilis]
https://smart.embl.de/smart/DDvec.cgi?smart=128:Pfam_Ribosomal_L17(16|128)+
1029190,24378360,50S ribosomal protein L17,AAN59603,AAN59603.1,GI:24378360,Large subunit ribosomal protein L17,NC_004350.2,NP_722297,NP_722297.1,Q8DS37,RL17_STRMU,Rl17,SMU_2000,WP_002262314.1,rl17,rplQ,smu:SMU_2000

rl29 210007.SMU_2019 Large subunit ribosomal protein l29; Belongs to the universal ribosomal protein uL29 family
https://smart.embl.de/smart/DDvec.cgi?smart=69:Pfam_Ribosomal_L29(11|66)+
1029208,24378380,50S ribosomal protein L29,50s ribosomal protein L29,AAN59622,AAN59622.1,GI:24378380,Large subunit ribosomal protein L29,NC_004350.2,NP_722316,NP_722316.1,Q8DS21,RL29_STRMU,Rl29,SMU_2019,WP_002262332.1,rl29,rpmC,smu:SMU_2019

rl31 210007.SMU_1298 Large subunit ribosomal protein l31; Belongs to the bacterial ribosomal protein bL31 family. Type B subfamily
https://smart.embl.de/smart/DDvec.cgi?smart=80:Pfam_Ribosomal_L31(1|78)+
1028520,24377673,50S ribosomal protein L31,50S ribosomal protein L31 type B,AAN58975,AAN58975.1,GI:24377673,Large subunit ribosomal protein L31,NC_004350.2,NP_721669,NP_721669.1,Q8DTN5,RL31B_STRMU,Rl31,SMU_1298,WP_002263148.1,rl31,rpmE2,smu:SMU_1298

rpoD 210007.SMU_822 Dna-dependent rna polymerase sigma subunit; Sigma factors are initiation factors that promote the attachment of RNA polymerase to specific initiation sites and are then released. This sigma factor is the primary sigma factor during exponential growth
https://smart.embl.de/smart/DDvec.cgi?smart=371:Pfam_Sigma70_r1_1(11|90)+Pfam_Sigma70_r1_2(99|132)+Pfam_Sigma70_r2(137|207)+Pfam_Sigma70_r3(216|293)+Pfam_Sigma70_r4(305|358)+ 1028204,24377195,AAN58538,AAN58538.1,DNA-dependent RNA polymerase sigma subunit,GI:24377195,NC_004350.2,NP_721232,NP_721232.1,O33662,RNA polymerase primary sigma factor,RNA polymerase sigma factor RpoD,RNA polymerase sigma factor SigA,RpoD,SIGA_STRMU,SMU_822,Sigma-42,WP_002261972.1,rpoD,sigA,smu:SMU_822

rpsD 210007.SMU_2135c Small subunit ribosomal protein s4; One of the primary rRNA binding proteins, it binds directly to 16S rRNA where it nucleates assembly of the body of the 30S subunit
https://smart.embl.de/smart/DDvec.cgi?smart=203:Ribosomal_S4(3|92)+S4(93|157)+
1029300,24378494,30S ribosomal protein S4,AAN59726,AAN59726.1,GI:24378494,NC_004350.2,NP_722420,NP_722420.1,P59133,RS4_STRMU,SMU_2135c,Small subunit ribosomal protein S4,WP_002262437.1,rpsD,smu:SMU_2135c
rpsI 210007.SMU_170 Small subunit ribosomal protein s9; Belongs to the universal ribosomal protein uS9 family
https://smart.embl.de/smart/DDvec.cgi?smart=130:Pfam_Ribosomal_S9(10|130)+
1029746,24376548,30S ribosomal protein S9,AAN57946,AAN57946.1,GI:24376548,NC_004350.2,NP_720640,NP_720640.1,Q8DW97,RS9_STRMU,SMU_170,Small subunit ribosomal protein S9,WP_002262992.1,rpsI,smu:SMU_170
scpA 210007.SMU_1713c Segregation and condensation protein a; Participates in chromosomal partition during cell division. May act via the formation of a condensin-like complex containing Smc and ScpB that pull DNA away from mid-cell into both cell halves
https://smart.embl.de/smart/DDvec.cgi?smart=235:Pfam_SMC_ScpA(18|231)+
1029544,24378082,AAN59348,AAN59348.1,Conserved hypothetical protein,GI:24378082,NC_004350.2,NP_722042,NP_722042.1,Q7ZAK8,SCPA_STRMU,SMU_1713c,Segregation and condensation protein A,WP_002262561.1,conserved hypothetical protein,scpA,smu:SMU_1713c
spaP 210007.SMU_610 Cell surface antigen spap; Surface protein antigen implicated in dental caries
https://smart.embl.de/smart/DDvec.cgi?smart=1562:Pfam_Strep_SA_rep(201|225)+Pfam_Strep_SA_rep(226|250)+Pfam_Strep_SA_rep(283|307)+Pfam_Strep_SA_rep(308|332)+Pfam_Strep_SA_rep(365|389)+Pfam_Strep_SA_rep(390|414)+Pfam_Strep_SA_rep(447|471)+Pfam_GbpC(492|816)+Pfam_Antigen_C(1328|1490)+Pfam_Gram_pos_anchor(1521|1562)+ 1028055,24376987,5LO3,5LO4,AAN58348,AAN58348.1,Cell surface antigen I,Cell surface antigen I/II,Cell surface antigen II,Cell surface antigen SpaP,GI:24376987,NC_004350.2,NP_721042,NP_721042.1,P23504,SMU_610,SPAP_STRMU,SpaP,WP_002352221.1,cell surface antigen SpaP,smu:SMU_610,spaP
tuf 210007.SMU_714 Translation elongation factor ef-tu; This protein promotes the GTP-dependent binding of aminoacyl- tRNA to the A-site of ribosomes during protein biosynthesis
https://smart.embl.de/smart/DDvec.cgi?smart=398:Pfam_GTP_EFTU(10|205)+Pfam_GTP_EFTU_D2(228|298)+Pfam_GTP_EFTU_D3(302|397)+
1028120,24377091,AAN58443,AAN58443.1,EF-Tu,EFTU_STRMU,Elongation factor Tu,GI:24377091,NC_004350.2,NP_721137,NP_721137.1,P72483,SMU_714,Translation elongation factor EF-Tu,WP_002263314.1,smu:SMU_714,translation elongation factor EF-Tu,tuf
